# Islet vascularization is regulated by primary endothelial cilia via VEGF-A dependent signaling

**DOI:** 10.1101/670760

**Authors:** Yan Xiong, M. Julia Scerbo, Anett Seelig, Francesco Volta, Nils O’Brien, Andrea Dicker, Daniela Padula, Heiko Lickert, Jantje M. Gerdes, Per-Olof Berggren

## Abstract

Accumulating evidence point to a role for primary cilia in endothelial cell function. Islet vascularization is an important determinant of islet function and glucose homeostasis. We have previously shown that β-cell cilia directly regulate insulin secretion. However, it is unclear whether primary cilia are also implicated in islet vascularization and thus contribute to glucose homeostasis.

**Objective:** To characterize the role of primary cilia in islet vascularization.

**Methods and Results:** At four weeks, *Bbs4^−/−^* islets show markedly lower intra-islet capillary density with enlarged diameters. We transplanted islets into the anterior chamber (ACE) of mouse eyes for longitudinal and non-invasive *in vivo* monitoring of vascular morphology. *Bbs4*−/− islets exhibited significantly delayed re-vascularization and enlarged vessels during engraftment. Similar vascular phenotypes were observed in two other ciliopathy models. By shifting the relative contributions of host versus donor endothelial cells in islet revascularization, we found that primary cilia on endothelial cells is essential for this process. Electron microscopy analysis further revealed a lack of fenestration in engrafted *Bbs4*−/− islets, partially impairing vascular permeability and glucose delivery to β-cells. Finally, we identified that Vascular endothelial cell growth factor A (VEGF-A)/VEGF receptor 2 (VEGFR2) signalling is involved in islet vascularization, islet function and vascular fenestration. *In vitro* silencing of two different ciliary genes in endothelial cells disrupts VEGF-A/ VEGFR2 internalization and phospho-activation of downstream signalling components. Consequently, key features of angiogenesis including proliferation, migration and tube formation are attenuated in *BBS4* silenced endothelial cells.

**Conclusions:** Endothelial cell primary cilia regulate islet vascularization and vascular barrier function via VEGF-A/ VEGFR2 signaling pathway. Islet vascularization is impaired in four weeks old *Bbs4^−/−^* mice. Long-time monitoring of re-vascularization of WT and *Bbs4^−/−^* islets recapitulates the phenotype and demonstrates a role for cilia in islet vascularization and vascular barrier function. VEGF-A/ VEGFR2-dependent signalling is regulated by endothelial primary cilia.

## Introduction

Organ growth, development and function are processes that are critically linked to the development of functional blood supply. Tissues rely on functional blood vessels for efficient delivery of oxygen or nutrients such as glucose. In addition, blood vessels are important to establish microenvironments or “niches” required for stem or progenitor cell maintenance among others^1^. Moreover, mis-regulated angiogenesis plays a role in numerous diseases, including but not limited to, diabetic complications, cancer progression and metastasis^2^. In the pancreas, the exocrine portion of the organ is organized by branching ducts while the endocrine islets of Langerhans are scattered throughout the exocrine tissue and interconnected by blood vessels. Pancreatic bud formation and blood vessel formation are initiated at the same time; however, although both endocrine and exocrine pancreatic tissue are derived from the same progenitors, vessel density in the exocrine portion of the organ is considerably lower compared to intra-islet vessel density. This suggests that separate factors play a role in endocrine pancreatic vascularization. In addition, islet re-vascularization has been suggested to be a critical determinant in graft survival following islet transplantation in the treatment of *Type 1 Diabetes*^4–6^.

One of the main players in islet vascularization is vascular endothelial growth factor A (VEGF-A)/ VEGF receptor 2 dependent signaling; VEGF-A is secreted by islet endocrine cells (both α- and β-cells) and interacts with VEGFR2 receptor on microvessels in the periphery to recruit blood vessels to the islets^5,7^. Constitutive, targeted deletion of *Vegfa* from β-cells revealed a critical role for VEGF-A dependent signals in blood vessel formation, maintenance and function^5^. Mice were severely glucose intolerant but showed normal islet function in isolated islets, suggesting that either nutrients (such as glucose) did not get delivered efficiently to the β-cells or insulin disposal into the blood stream was significantly hampered. In contrast, removal of *Vegfa* from mature β-cells had a less severe impact on glucose tolerance although intra-islet capillary density was reduced by 50%^8^; this could suggest that, after islet maturation, local VEGF-A/ VEGFR2 dependent signaling is less important for the maintenance of intra-islet capillaries and insulin disposal compared to during development.

Primary cilia are present on roughly eighty percent of the cells in the adult body plan^9^. Among others, endothelial cells are ciliated and endothelial cilia have been implicated in flow-sensing and vascular hypertension, intracranial blood vessel formation, and atherosclerosis prevention^10–12^. Recent studies also unveiled a novel role of primary cilium in preventing vascular regression^13^. In metabolically active organs, they play a role in sensing metabolic signals and energy homeostasis^14 15^. In addition, we have shown that β-cell cilia play a role in insulin signaling, insulin secretion and glucose homeostasis^16^ (Volta et al, *in revision).* Endothelial cells in both central and peripheral organs of insulin action have been implicated in insulin resistance^17^ and diabetes etiology^18^. Here, we present one of the first studies addressing the role of endothelial cilia in islet vascularization and re-vascularization of transplanted islets.

## Results

### Intra-islet capillary density is reduced in Bbs4^−/−^ pancreata

To test if primary cilia play a role in pancreatic islet vascularization, we characterized intraislet capillaries in pancreatic cryosections of two-month old *Bbs4^−/−^* mice (N=8). Platelet endothelial cell adhesion molecule (PECAM-1) immunofluorescence as a marker of endothelial cells revealed 35% reduced intra-islet capillary density compared to wildtype (wt) littermate controls (Fig. 1A, P=0.0019). To calculate intra-islet capillary density, the relative PECAM-1 positive volume was normalized to insulin positive islet volume (Fig. 1B). In addition, the average vessel diameter was increased by 18% compared to that in wt controls (Fig. 1C, P<0.0001). Because tissue integrity is often compromised in cryopreserved samples, we corroborated our results by whole mount staining and imaging of freshly isolated islets of two-month old mice (Fig. 1D-1F). In good agreement with our previous observations, capillary density was reduced by 33% and vessel diameter increased by 18% in islets of *Bbs4^−/−^* mice (P=0.0044 and 0.0067 respectively). Of note, we observed no change in pericyte coverage of intra-islet capillaries based on Neuron-glial 2 (NG2) immunofluorescence (Suppl. Fig. 1A, 1B, P=0.7045). In four-month old *Bbs4^−/−^* mice, however, the difference in intra-islet capillary density and diameter in *Bbs4^−/−^* mice approximated that of wt littermates (Supp. Fig. 1C-1E, P=0.0676 and 0.0736 respectively). These dynamics implicate that primary cilia and centrosomal/ basal body integrity play a role during pancreas and islet development. Importantly, there was no detectable difference in vascularization of the exocrine portion of the pancreas at four months of age (Suppl. Fig. 1F). Therefore, *Bbs4* function seems to be mostly restricted to the endocrine pancreas and not relevant for the exocrine compartment.

**Fig 1.**
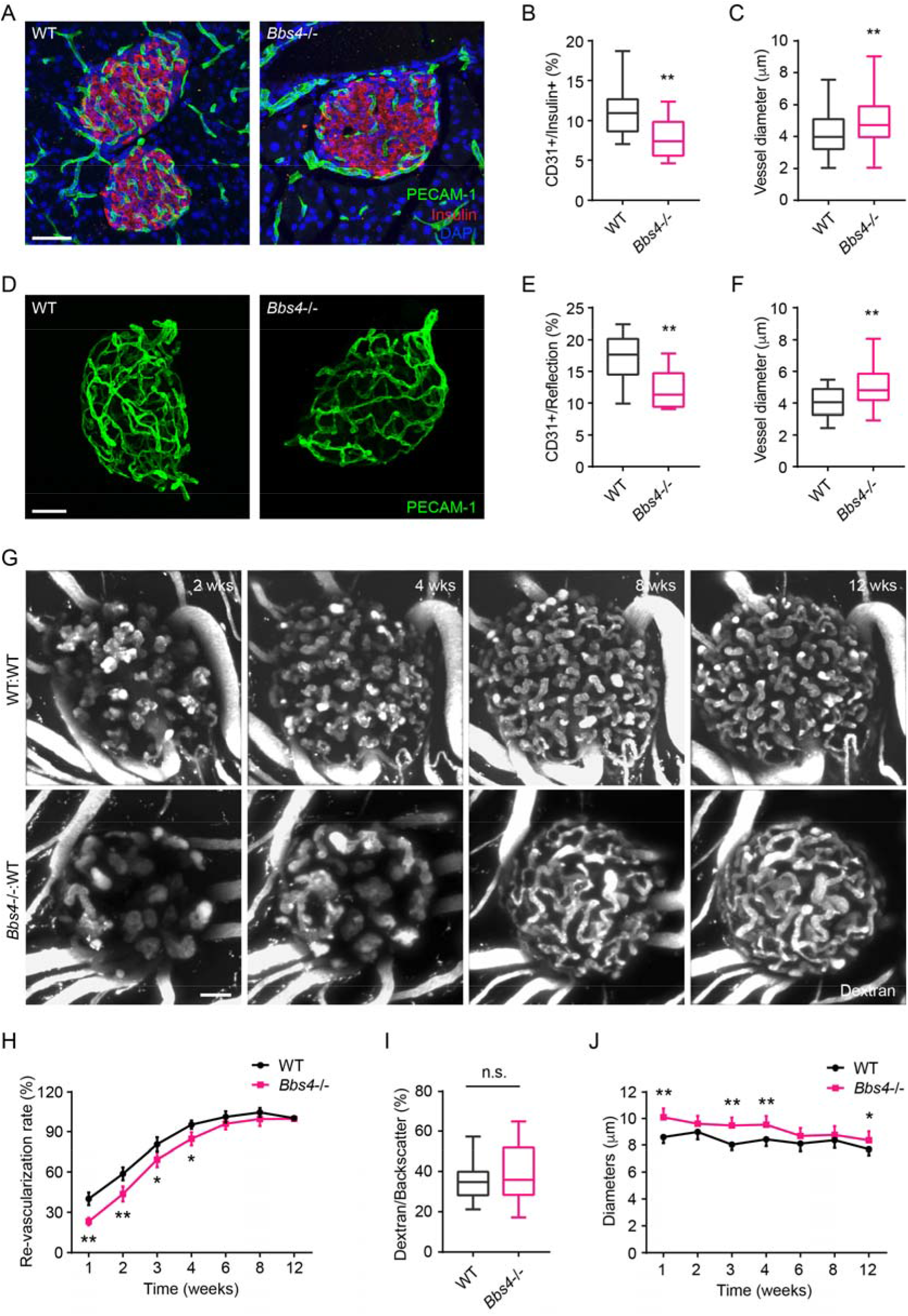
*Bbs4−/−* islets show delayed vascularization and enlarged capillary diameter in the pancreas and transplantation site. **(A)** Immuno-fluorescence staining of pancreatic sections from 2-month-old wt and *Bbs4−/−* mice showing islets (insulin, red) and intra-islet capillaries (PECAM-1, green). **(B-C)** Quantification of relative intra-islet PECAM-1 positive volume, normalized to insulin positive volume (B) and average intra-islet capillary diameters (C) in wt and *Bbs4−/−* pancreatic sections. Results are shown as box-and-whisker plots (running from minimal to maximal values), **p<0.01, n=8 for animals and n=10 islets per animal. **(D)** Immunofluorescence staining of freshly isolated and fixed pancreatic islets from 2-month-old wt and *Bbs4*−/− mice, showing PECAM-1 (green) labelled islet capillaries. **(E-F)** Quantification of relative CD31 positive volume within each islet normalized to islet volume estimated by backscatter signal (E) and average capillary diameters (F) in wt and *Bbs4−/−* islets. Results are shown as box-and-whisker plots (running from minimal to maximal values), **p<0.01 by t-test, n=3 for animals and n=4 islets per animal. **(G)** Re-vascularization of 2 day-cultivated wt (upper) and *Bbs4−/−* (lower) islets in wt recipient eyes at 2, 4, 8- and 12-weeks post transplantation, visualized by intravenous injection of Texas Red-conjugated dextran. **(H)** Quantification of re-vascularization rates of wt and *Bbs4*−/− islet grafts in wt recipients. Results are mean±S.E.M. **(I)** Relative vascular density of wt and *Bbs4−/−* islet grafts at the end of 12 weeks post transplantation. Results are shown as box-and-whisker plots (running from minimal to maximal values), n.s. means not significant by t-test. **(J)** Average diameters of newly formed capillaries in wt and *Bbs4−/−* islet grafts in wt recipients. Results are mean±S.E.M. \**p*<0.05, \*\**p*<0.01 by two-way-ANOVA, n=6 for animals and n=5 islets per animal. Scale bars: 50 μm.

### Bbs4^−/−^ endothelial cells exhibits blunted angiogenic response during islet re-vascularization

To test if cilia play a role in intra-islet capillary formation during engraftment of transplanted tissue as well as during development, we transplanted murine islets into the anterior chamber of the eye (ACE)^19^. This approach allows for longitudinal and non-invasive *in vivo* monitoring of islet engraftment and re-vascularization. We and others have previously shown that endothelial cells disappear over prolonged periods of cultivation of isolated islets. Immediately after isolation, endothelial cells maintain the intra-islet capillary network. After two days in culture, endothelial cell clusters remain. Seven days post isolation endothelial cells have disappeared from the islet (Suppl. Fig. 2A). Thus, when transplanted shortly after isolation, a significant number of endothelial cells from the donor survives and contributes to revascularization of the islet graft^4,20,21^. Importantly, both islet cells and endothelial cells are ciliated. Therefore, to determine whether the role of primary ciliary/ centrosomal function in islet endothelial or endocrine cells is underlying the reduction in intra-islet capillary density, we capitalized on the temporal differences in the ratio of donor or recipient endothelial cells in intra-islet capillaries (Suppl. Fig. 2B).

In one experimental setting, islets isolated from *Bbs4^−/−^* and wt littermate controls were cultivated for two days before transplantation into the ACE of wt recipients (Fig. 1G, Suppl. Fig. 2B, upper panel). We determined the rate of re-vascularization by intravenous (iv) injection of fluorescently labeled dextran once a week for a total of twelve weeks (Fig. 1G). The percentage of dextran-related fluorescence in total islet volume (based on backscatter signal^22^) was used as a measure of islet vascularity, and re-vascularization rate of each islet was calculated by normalizing its dextran-related volume at each individual week to the volume at twelve weeks. Two weeks after transplantation, *Bbs4^−/−^* islets showed significantly less re-vascularization compared to wt controls (Fig. 1H). Until four weeks after transplantation, re-vascularization remains lower than that of wt islets (Fig. 1H). Twelve weeks post transplantation, intra-islet vascular density measured as dextran related fluorescence intensity relative to islet volume is similar in both *Bbs4^−/−^* and wt islets (Fig. 1I). Morphologically, the newly formed blood vessels vary among the two different genotypes: the vessel diameter is greater in *Bbs4^−/−^* islets, suggesting that there are fewer, wider islet capillaries in these islets compared to the wt controls during the first four weeks of engraftment (Fig. 1J), similar to the results obtained from cryosections of young *Bbs4^−/−^* animals.

In the other experimental setting, islets were kept in culture for seven days after isolation before transplanting to the ACE of wt recipients; this treatment ablates endothelial cells in the donor tissue and favors almost exclusively revascularization by the recipients’ vascular system (Suppl. Fig 2B, lower panel). We did not observe significant differences in re-vascularization rate over time or islet vascular density at the twelve-week endpoint (Fig. 2A-2C). In addition, vessel morphology and diameter were similar in *Bbs4^−/−^* and wt control islets (Fig. 2D). Because *Bbs4^−/−^* endothelial cells were lost over the seven days cultivation period, our findings suggest that primary cilia on endothelial cells are required for the angiogenic response and thus islet re-vascularization.

**Fig 2.**
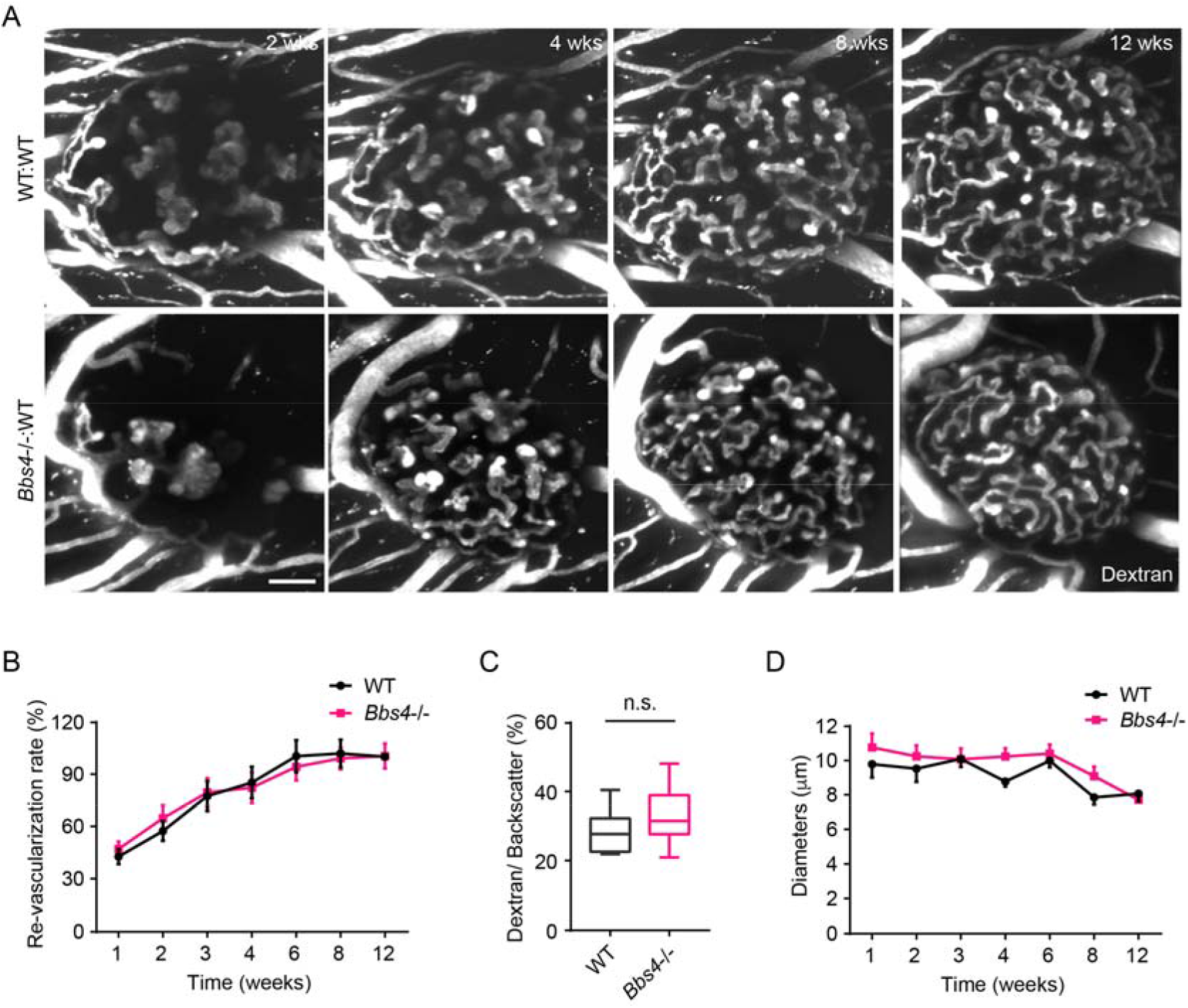
*Bbs4*−/− islets after prolonged culture exhibits normal re-vascularization patterns in wt recipient eyes. **(A)** Re-vascularization of 7 day-cultivated wt (upper) and *Bbs4*−/− (lower) islets in wt recipient eyes at 2, 4, 8- and 12-weeks post transplantation, visualized by intravenous injection of Texas Red-conjugated dextran. Donor islets have been cultured for 7 days prior to transplantation. **(B)** Quantification of re-vascularization rates of wt and *Bbs4−/−* islet grafts in wt recipients. Results are mean±S.E.M. **(C)** Relative vascular density of wt and *Bbs4−/−* islet grafts at the end of 12 weeks post transplantation. Results are shown as box-and-whisker plots (running from minimal to maximal values), n.s. means not significant by t-test. **(D)** Average diameters of newly formed capillaries in wt and *Bbs4−/−* islet grafts in wt recipients. Results are mean±S.E.M., n=4 for animals and n=5 islets per animal. Scale bar: 50 μm.

To verify our conclusion, we reversed the transplantation strategy by transplanting wt islets to *Bbs4^−/−^* recipients either directly after isolation or after keeping them in culture for two and seven days respectively (Suppl. Fig. 3). To avoid confounding effects of glucose intolerance on islet engraftment, we transplanted islets into the eyes of four months old *Bbs4^−/−^* mice that were obese but not diabetic yet (Suppl. Fig. 3A-3C). Based on the apparent role of endothelial primary cilia in the angiogenic response, we expected stronger impairment in re-vascularization compared to wt recipients. Indeed, we observed significantly slower revascularization in this transplantation scheme and the rates correlated with the time islets were kept in culture (Suppl. Fig. 3D, 3E). The revascularization rate of wt islets transplanted to the ACE of *Bbs4^−/−^* mice seven days post-isolation was significantly lower than that of wt islets two days post-isolation (P=0.0147), and both were significantly lower than that of wt islets transplanted to wt ACE (P=0.0111). Interestingly, intra-islet vessel density after completion of re-vascularization was not significantly lower than that of wt islets transplanted into wt recipients at the twelve week endpoint (Suppl. Fig. 3F). The diameter of intra-islet capillaries, however, was markedly wider in wt islets transplanted to *Bbs4^−/−^* recipients than the other two groups after seven days in culture (Suppl. Fig. 3G).

### Primary cilia of endothelial cells regulate islet re-vascularization

Because Bbs4 protein might have additional roles in cellular processes unrelated to the basal body, we used two additional mutant mouse models. *Pitchfork* (*Pifo*^−/−^) mice have dysfunctional cilia that cannot be properly disassembled^22^. We determined intra-islet vascular density in three-month old *Pofo^−/−^* mice; PECAM-1 staining revealed 20% reduction in intraislet vascular density and 30% increase in vessel diameter in *Pifo*^−/−^ mice compared to littermate controls, similar to our observation in *Bbs4^−/−^* islets (Suppl. Fig. 4A-4C, P=0.0421 and P=0.0032 respectively). Transplantation of *Pifo^−/−^* islets into wt ACE showed a delay in relative re-vascularization compared to wt islets, also similar to what we observed for *Bbs4^−/−^* mice (Suppl. Fig. 4D-4G). Although the effects on re-vascularization were less severe than those in *Bbs4^−/−^* islets, the findings suggest a role for primary cilia in islet vascularization and the angiogenic response in islet re-engraftment. To better understand the contributions of β-cell vs. endothelial cilia and confirm the findings from *Bbs4^−/−^* mice, we specifically ablated *Intraflagellar transport 88 (Ift88),* a core ciliary protein, in β-cells by crossing placing a floxed *Ift88* line to an inducible cyclic recombinase under the control of the *Pdx1-* promoter. This generates a β-cell specific, inducible ciliary knockout mouse (*Ift88*^loxP/loxP^; *Pdx1-CreER*) with an efficiency of 50%, that we will refer to as βICKO (pronounced BICKO) from here on (Volta et al., *in revision*) (Suppl. Fig. 5A). Recombination was induced between postnatal day 25 and 35 and we quantified intra-islet vascular density in Tx-treated βICKO mice six weeks after induction; we found no observable difference between induced βICKO mice and oil-treated controls (Suppl. Fig. 5B, 5C). In addition, we isolated βICKO islets two months after Tamoxifen treatment and transplanted them into wt mice after overnight culture. We did not observe differences in relative re-vascularization rate between wt and *Ift88^Δ/Δ^; Pdx1-CreER* islets (Suppl. Fig. 5D-5G). Overall, our observations strongly suggest that cilia in endothelial cells regulate re-vascularization during islet engraftment.

### Intra-islet vasculature plays a role in glucose metabolism

Besides a significant higher vascular density than exocrine pancreas, another hallmark of intra-islet capillary is fenestrae as interfaces for substance exchange between islets and the blood stream^23^. Thus, we further examined the barrier function of *Bbs4^−/−^* islets four months after engraftment, when the intra-islet vascular density is already comparable with wt. As previously reported, in isolated islets or *Bbs4^−/−^* depleted MIN6m9 cells, we did not observe a difference in β-cell glucose uptake or glucose metabolism^16^. However, glucose and insulin diffusion through of *Bbs4^−/−^* depleted islet endothelial cells remain uncharacterized. Electron micrographs revealed significantly reduced fenestration in the capillaries of *Bbs4*^−/−^ compared to wt islet grafts in the ACE four months after transplantation (Fig. 3A, 3B, 27% lower, P=0.0006). Although the basement membrane seemed slightly thicker in *Bbs4^−/−^* islets, we did not observe significant thickening of this structure (Fig. 3C, P=0.0571). To test if reduced fenestration could affect the permeability and barrier function of *Bbs4^−/−^* islet capillaries, we injected 40 kDa fluorescently labeled dextran and measured the diffusion of dextran-related fluorescence outside the islet graft over time (Fig. 3D). During the time of the experiments, we observed an immediate increase of fluorescence intensity outside the islet graft followed by a plateau that was established at comparable times after dextran addition due to the dynamic aqueous humor flow. Importantly, both the initial rate and the plateau were decreased in *Bbs4^−/−^* islet grafts (Fig. 3E, P=0.0437). Simulation of this process also revealed different kinetics of dye leakage (Fig. 3F), suggesting that the delivery of medium sized molecules is less efficient in *Bbs4^−/−^* intra-islet capillaries. Nutrients such as glucose or fatty acids are small molecules often relying on active transport across cell membranes. But in the case of highly fenestrated capillaries, solutes including small molecules, ions and hormones pass freely through the pores between the blood and abluminal surface. To test if glucose delivery to β-cells were affected by the ultrastructural changes in islet capillaries, we injected fluorescently labeled glucose analog (2-NBDG) into animals transplanted with *Bbs4^−/−^* or wt islets. 2-NBDG was rapidly transported into β-cells from intra-islet capillaries and simultaneously leaked into the surrounding aqueous humor from iris vessels. We followed glucose uptake by measuring fluorescence intensity originating from β-cells and normalized it to the fluorescence intensity originating from the aqueous humor over time (Fig. 3G). Importantly, wt β-cells internalized significantly more 2-NBDG than *Bbs4^−/−^* β-cells under the same period of time (Fig. 3H, P=0.0418). This result points to a critical role of ciliary/ basal body proteins in maintaining islet capillary fenestration with implications for the delivery of glucose and potentially other nutrients to islet cells. To investigate the physiological relevance of our finding, we utilized our transplantation model in combination with *Bbs4^−/−^* or wt islets to better understand how reduced intra-islet capillary density and permeability may affect islet output and glucose tolerance. It was previously shown that 100 islets (diameter between 150 and 200 μm) transplanted into the ACE of chemically induced diabetic mice were sufficient to revert blood glucose levels to normoglycemic levels; this amount of pancreatic islets is defined as marginal islet mass^24^. C57/B6 J mice were treated with strepotozotocin and subsequently transplanted with marginal islet mass of wt or *Bbs4^−/−^* islets. Four weeks after transplantation, mice transplanted with *Bbs4^−/−^* islets were significantly glucose intolerant compared to those mice transplanted with wt islets (Fig. 3I). Eight weeks after transplantation, mice showed impaired glucose tolerance and, 12 weeks posttransplantation, only mildly impaired glucose handling compared to those transplanted with wt islets (Fig. 3J, 3K). Overall, the phenotype is ameliorated over time, which is consistent with our previous observations in re-vascularization (Fig. 1, 2), suggesting that a lack of capillaries in *Bbs4^−/−^* islets indeed negatively affects whole body glucose homeostasis. Glucose handling is mildly impaired by the end of 12 weeks, which is probably caused by impaired glucose delivery to β-cells and a combination of defective 1^st^ phase insulin secretion^16^.

**Fig 3.**
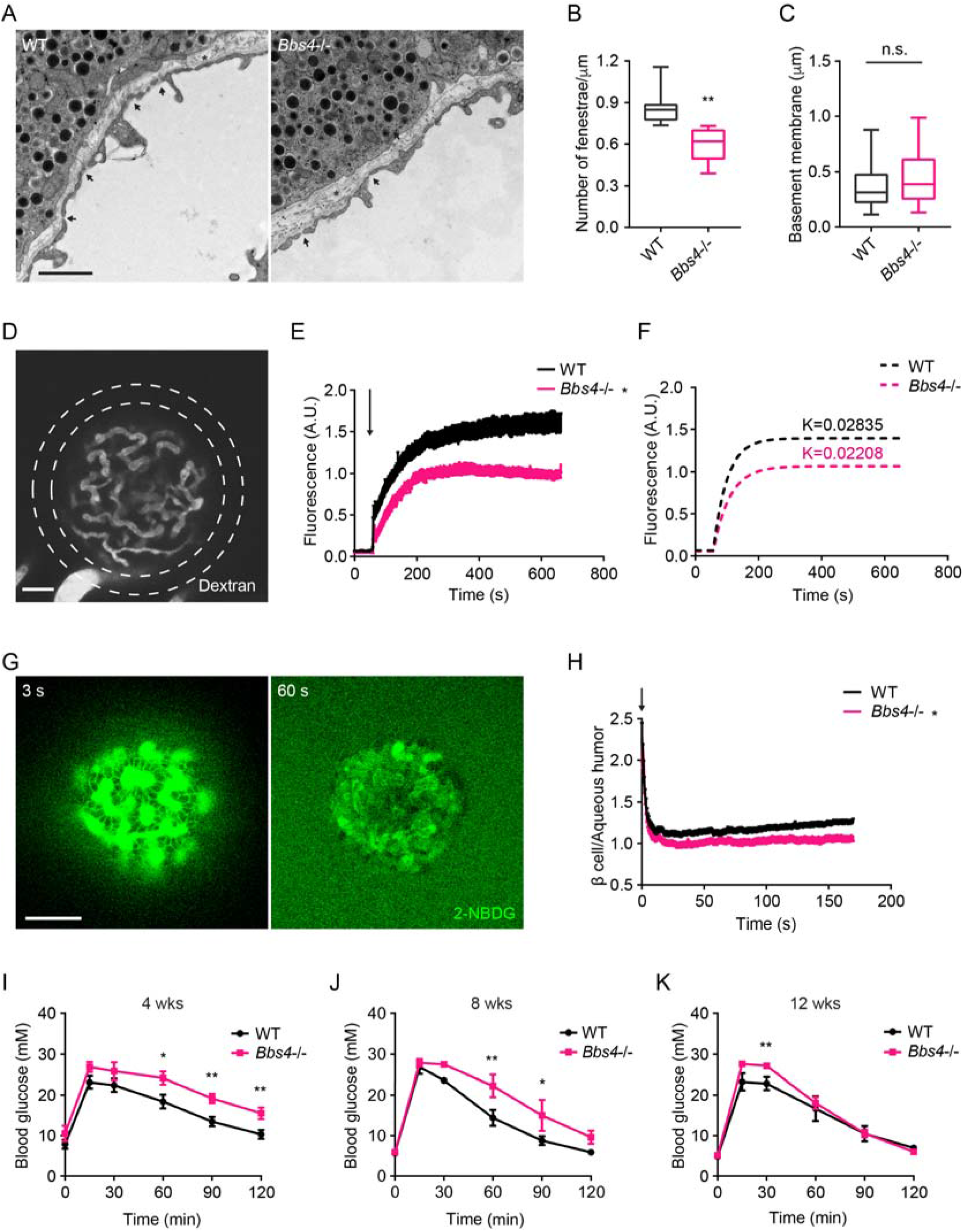
Dysfunctional intra-islet vasculature in *Bbs4−/−* islets undermines glucose metabolism **(A)** Electron microscopic images of wt and *Bbs4−/−islet* graft dissected from wt recipient eyes, showing fenestrated islet capillaries (arrows) and basement membrane (asterisks). Scale bar: 1 μm. **(B-C)** Quantification of average fenestrae density (B) and basement membrane thickness (C) of the capillaries in wt and *Bbs4−/−* islet grafts. Results are shown as box-and-whisker plots (running from minimal to maximal values), \*\**p*<0.01, n.s. means not significant by t-test, n=7. **(D)** Representative image showing leakage of 40 KDa FITC conjugated dextran from wt islet grafts in mouse eyes at 1 min after injection. Scale bar: 50 μm. **(E-F)** Quantification of FITC fluorescence intensity in the region circles by the dashed lines outside wt and *Bbs4−/−*islet grafts (E) and simulated curves showing different kinetics **(F)**. Arrow indicates injection time point. Results are mean±S.E.M. *p<0.05 by two-way-ANOVA, n=12 for animals. (G) Representative image showing 2-NBDG leakage from wt islet vasculature and uptake by β cell in vivo. Times points are 3 s (left) and 60 s (right) after injection. Scale bar: 50 μm. (H) Real time ratio of 2-NBDG fluorescence intensity in β cells of wt and *Bbs4−/−* islet grafts to aqueous humor. Arrow indicates injection time point. Results are mean± S.E.M. \**p*<0.05, n=8 for animals. **(I-K)** Intraperitoneal glucose tolerance test of wt recipient mice which were transplanted with WT or *Bbs4*−/− islets, at 4 weeks (I), 8 weeks (J) and 12 weeks (K) post transplantation. Results are mean±s.e.m. \**p*<0.05, \*\**p*<0.01 by two-way-ANOVA, n=6 for animals.

### *Impaired VEGFA-VEGFR2 dependent signaling in* Bbs4-*depleted endothelial cells*

Intra-islet capillary density and fenestration is primarily maintained by VEGF-VEGFR2-signaling^25–27^. To test the expression levels of VEGF-A and other key angiogenic factors players in Bbs4-depleted endothelial cells, we suppressed *Bbs4* by RNA interference in a pancreatic endothelial cell line, MS1. We chose the shRNA which reduced *Bbs4* levels by two-fold, and examined the expression of the same set of angiogenesis regulators (Suppl. Fig. 6A). We found no observable change in Tie1, Tie2, endothelial Nitric oxide synthase eNOS or Notch ligand Delta-4 (Dll4), while VEGFR2, Ephrin A1 and B2 were downregulated by 20% (Fig. 4A).

**Fig 4.**
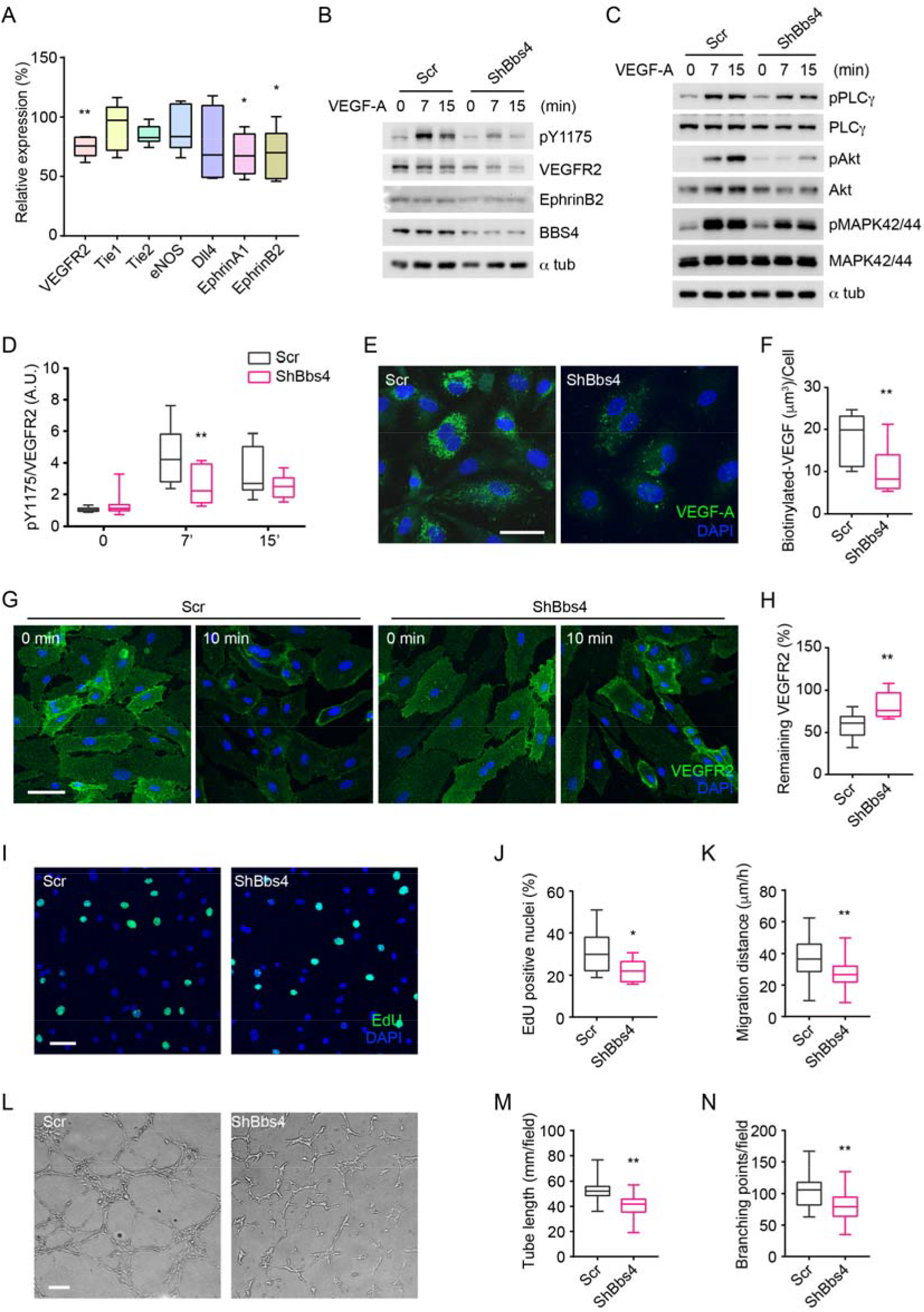
VEGF-VEGFR2 induced signalling pathway and angiogenic response is disrupted in *Bbs4* silenced endothelial cells. **(A)** qPCR analysis of VEGF signaling related gene expression in *Bbs4* silenced MS1. Results are presented as relative mRNA levels normalized to scrambled shRNA treated cells. Data is shown as box-and-whisker plots (running from minimal to maximal values), \**p*<0.05, \*\**p*<0.01 by t-test, n=3. **(B)** and **(C)** Western blots showing VEGF signaling pathway in scrambled or shBbs4 treated HDMEC. **(D)** Quantification of VEGFR2 activation. **(E)** Representative images showing biotinylated VEGF-A uptake in scrambled or ShBbs4 treated endothelial cells and quantification **(F). (G)** Representative cell surface staining of VEGFR2 in scrambled (Scr) or ShBbs4 treated HDMECs, prior to (0 min) and after VEGF stimulation (10 min). Scale bar: 50 μm. **(H)** Quantification of the percentage of remaining membrane-bound VEGFR2. **(I)** VEGF induced proliferation in Scr or ShBbs4 treated MS1 by EdU assay (green) and quantification **(J)**. Scale bar: 50 μm. **(K)** VEGF induced endothelial migration in Scr or ShBbs4 treated MS1. **(L)** VEGF induced endothelial matrigel tube formation in Scr or ShBbs4 treated MS1, and quantification of total tube length **(M)** and branching points **(N)**. Scale bar: 200 μm. Results are shown as box-and-whisker plots (running from minimal to maximal values), \**p*<0.05, \*\**p*<0.01 by t-test, n=3.

Angiogenesis is not exclusively regulated on the transcriptional level but also relies on a complex network of signaling cascades. Therefore, we tested if VEGFR2-dependent phospho-activation of downstream signaling molecules, including Phospholipase C*γ*1 (PLC*γ*1), Protein Kinase B (Akt) and Mitogen-activated protein kinase (MAPK 42/44) were affected. We used HDMEC cells stably transfected with shRNA targeting *BBS4* and tested before, seven and fifteen minutes after VEGF-A addition (Fig. 4B–4D, Suppl Fig. 6A, 6B). In control cells (scrambled), VEGFR2 Tyr 1175 phosphorylation is markedly increased at seven minutes (4.3 fold) and sustained until 15 minutes (3.4 fold) after stimulation (Fig. 4B, 4D). By comparison, pTyr1175 levels increased minimally in BBS4-depleted HDMEC cells (2 fold and 1.8 fold respectively, Fig. 4B, 4D). At the same time, total levels of VEGFR2 are also lowered in absence of Bbs4 by 20% (Fig. 4B, Suppl. Fig. 6B, P=0.0101). In addition, we found that EphrinB2 protein level is slightly reduced in BBS4-depleted HDMEC (Fig. 4B, Suppl. Fig. 6B, 15%, P=0.0051), corroborating our previous finding of decreased gene expression (Fig. 4A). While PLC*γ*1 phosphorylation peaks at seven minutes and decreases 15 minutes after stimulation in control cells, loss of Bbs4 leads to a 40% attenuation in PLC*γ*1 phosphorylation at seven minutes (Fig. 4C, Suppl 6C, P=0.0129). Similarly, pAkt and pMAPK 42/44 were significantly reduced in BBS4-depleted cells seven and 15 minutes after VEGF-A addition (Fig. 4C, Suppl 6C, P=0.044 and P=0.0083 respectively). We further validated these results in isolated islets from wt and *Bbs4*^−/−^ animals. In unstimulated islet cells, pMAPK levels did not differ between wt and *Bbs4*^−/−^ cells. Ten minutes after addition of 100 ng/ml VEGF-A, pAkt levels increased 2.2-fold in wt but only 1.4-fold in *Bbs4^−/−^* cells. Similarly, pMAPK levels increased 3-fold in wt but only 1.5-fold in *Bbs4^−/−^* cells, demonstrating that VEGF-A triggered downstream signaling is not sufficiently activated in these cells (Suppl. Fig. 6D, 6E).

Binding of VEGF-A to receptors triggers their endocytosis instantly, which is the initial step for subsequent intracellular signaling, and mainly regulated by EphrinB2^28–30^. Several studies have shown that VEGFR2 does not signal from the plasma membrane but from other intracellular compartments, mainly early endosomes and the endosomal sorting complex required for transport (ESCRT)^30,31^. The observed attenuation of VEGFR2 downstream signaling may stem from dysregulation of these initial steps. Therefore, we examined the uptake of biotinylated VEGF-A in BBS4-depleted HDMECs. After 30 min incubation, BBS4-depleted cells had only internalized 50 % VEGF-A compared to controls (Fig. 4E, 4F). We also tested VEGF-A uptake dynamics in a pancreatic endothelial cell line, MS-1 cells. Similar to our observations in HDMEC, control MS-1 cells efficiently and rapidly internalized VEGF-A, while by comparison, Ift88-depleted MS-1 cells internalized less VEGF-A after 30 min (58% compared to controls, Suppl Fig. 6F-6H). On the other hand, we also tested if VEGFR2 internalization was impaired in primary human endothelial cells depleted of *BBS4* (Fig. 4G). Plasma membrane bound VEGFR2 was evident prior to VEGF-A stimulation in both control and BBS4-depleted HDMEC cells. After ten min incubation with VEGF-A, a significant number of VEGFR2 was internalized in control cells, while 25% more VEGFR2 remained on the cell in BBS4-silenced cells, which indicates decreased internalization of VEGFR2 (Fig. 4H) and may be explained by our previous finding of lowered EphrinB2 levels (Fig. 4A, 4B). These results suggest that the VEGFR2 internalization upon ligand binding is regulated by ciliary and basal body proteins.

Furthermore, we examined the relative expression of the same set of angiogenic factors in isolated islets and found most expression levels unchanged, while VEGF-A expression was reduced by 20% in *Bbs4^−/−^* islets compared to wt controls. *Von-Hippel-Landau* gene expression, a known ciliary protein mutated in Von Hippel-Landau disease and a known regulator of Hypoxia induced factor 1a, was also unchanged (Suppl. Fig. 6I). VEGF-A gene expression was also slightly decreased in βICKO islets (Suppl. Fig. 6J). To test if this reduction was physiologically relevant, we tested VEGF-A secretion in *Bbs4^−/−^* and βICKO islets. Loss of Bbs4 or Ift88 function respectively decreased VEGF-A secretion by 33% and 28% (Suppl. Fig. 6K, P=0.0063 and Suppl. Fig. 6L, P=0.007), which could be explained by the reduction in VEGF-A gene expression. Altogether, these results consistently suggest that the intra-islet VEGF-A signaling pathway is impaired in *Bbs4^−/−^* islets.

VEGF-A signaling triggers a complex array of biological processes including endothelial cell proliferation and migration, two characteristic processes of angiogenesis. We tested the effect of BBS4-depletion on proliferation by ethynyldeoxyuridine (EdU) incorporation in MS1. Under the chosen conditions, VEGF-A induced cell proliferation in BBS4-depleted cells was reduced by 40% compared to controls cells (Fig. 4I, 4J, P<0.0001). A scratch assay revealed that BBS4-depleted cells covered only 73% of the distance of control cells in the same time frame (Fig. 4K, P<0.0001), similar to previous reports of ciliary involvement in cell migration^13^. Matrigel tube formation is a common in vitro assay for angiogenesis and we found that Bbs4-depleted cells formed less organized tubules and a subset of cells was not incorporated into tubular structures at all (Fig. 4L). Total tubular length was 80% in Bbs4-depleted compared to control cells (Fig. 4M, P<0.0001)). In addition, tubular structures formed by Bbs4-depleted cells established significantly fewer branching points than control cells (Fig. 4N, 75%, P<0.0001). These independent lines of evidence – reduced endothelial cell proliferation, migration and tube formation, and impaired vessel fenestration – all point to a role for cilia and basal body proteins in VEGF-A/ VEGFR2-dependent regulation of endothelial cell function.

## Discussion

Several recent studies have addressed the role of primary cilia in endothelial function^10–13^. However, the precise role of primary cilia in islet vascularization remains unclear. Here, we present one of the first studies revealing the role of the ciliary/ basal body machinery in islet vascularization both during islet maturation, adult tissue homeogenesis and islet re-vascularization.

*Bbs4^−/−^* islets exhibited lower intra-islet capillary density and enlarged capillaries at two months of age. Similarly, *Pifo^−/−^* islets also have a reduced intra-islet capillary density and increased vessel diameter compared to littermate controls at the same age. This suggests that the observed defects are probably due to dysfunctional cilia and not related to other, non-ciliary functions of *Bbs4*.

Islet vascularization is typically regulated by paracrine signaling between endocrine cells (including α- and β-cells) and peripheral endothelial cells. Islet transplantation into the ACE and subsequent non-invasive and longitudinal monitoring of re-vascularization recapitulated this phenotype. Several lines of evidence suggest that the observed phenotype is largely driven by impaired ciliary and basal body function in endothelial cells: first, *Bbs4^−/−^* islets that were transplanted two days after isolation were re-vascularized at a lower rate than those transplanted after seven days. Prolonged islet cultivation shifts the relative contribution to islet re-vascularization from *Bbs4^−/−^* depleted (donor) to wt (recipient) cells. Second, wt islets transplanted into the ACE of *Bbs4^−/−^* mice were more slowly re-vascularized when transplanted after seven days compared to two days of tissue culture. These conditions shift the composition of the islet vasculature toward a higher proportion of ciliary/ basal body deficient *Bbs4*^−/−^ endothelial cells. Third, native and transplanted βICKO islets (β-cell specific *Ift88* depletion) show no obvious defect in capillary density, diameter or revascularization rate. Although we cannot exclude a minor contribution by reduced VEGF-A secretion from both *Bbs4^−/−^* and βICKO islets, the effect on islet vascularization is likely negligible and mostly compensated by circulating VEGF-A, and we suggest there is an important role for the endothelial ciliary/ basal body apparatus in islet vascularization.

VEGF-A/ VEGFR2 is a key signaling pathway for maintaining the especially dense and leaky islet microvasculature. Our results suggest that dysfunctional primary cilia alter the VEGF-A/ VEGFR2 paracrine signaling activity in *Bbs4^−/−^* islets by impairing internalization of ligand-bound VEGFR2 receptor. VEGF-A/ VEGFR2 internalization and subsequent phosphorylation is disrupted in endothelial cells depleted of *BBS4* and *Ift88* respectively. Several downstream signaling components are insufficiently activated in good agreement with previous observations. We conclude that the observed phenotypes of reduced intra-islet capillary density and impaired fenestration of intra-islet micro-vessels are caused by defective VEGF-A/ VEGFR2 dependent signaling in endothelial cells.

Intra-islet capillary networks provide endocrine cells with nutrients, oxygen and growth factors which support the development and maturation of islets. In addition, the micro vasculature constantly adapts its barrier properties to the dynamic metabolic rates^32^. Finely tuned interplay between nutrient delivery to islet endocrine cells, endocrine cell function and insulin disposal into the blood stream is essential for the maintainance of glucose homeostasis^5,33^. Whereas intra-islet capillary density normalizes over time in *Bbs4^−/−^* islets, vessel function remains impaired when cilia/ basal function is compromised more than four months after transplantation. Blood vessel fenestration is significantly reduced in *Bbs4^−/−^* islets. Fenestrae are key features of islet capillaries that enable efficient nutrient delivery from and insulin disposal into blood vessels. *In vivo* 2-NBDG diffusion is blunted and leakage of 40 KDa dextran diminished when *Bbs4^−/−^* is depleted from intra-islet capillaries. With respect to glucose handling, decreased vessel permeability in *Bbs4*^−/−^ islets could impose an additional barrier on insulin disposal, adding to the observed defects in 1^st^ phase insulin. We thus suggest that defective delivery of nutrients such as glucose to β-cells and the disposal of insulin from β-cells into the blood stream may both contribute to impaired glucose handling of *Bbs4^−/−^* mice^16^.

Functional imaging of micro-vessels remains challenging and is largely limited to easily accessible tissues such as cornea and skin^34,35^. Transplantation of islets into the ACE not only renders established intra-islet capillaries more accessible for standard imaging techniques; it also provides the opportunity to follow dynamic processes such as islet revascularization under physiological as well as pathological conditions. This technique could also prove to be a useful resource for further investigation into blood vessel function in other tissues when transplanted into the ACE.

Finally, re-vascularization and establishing a functional interface between islet cells and blood supply are not only important in the context of islet transplantation but will likely prove vital for all approaches to β-cell replacement therapy. Moreover, taking into account the scarcity of organ donations worldwide, it is of paramount importance to optimize the outcomes of organ transplants. Revascularization rates in liver transplants correlate with long term graft survival, renal function and early allograft dysfunction^36^. Understanding the signaling role of primary cilium in this process provides us with novel targets to improve tissue re-vascularization and hence graft survival.

## Author contributions

Y.X and J.M.G designed the research and wrote the manuscript. Y.X, M.J.S, A.S, F.V, N.O’B, A.D performed the experiments. Y.X analyzed the data. J.M.G. had the idea for the study, D.P. and H.L. contributed the *Pifo^−/−^* mice, P.O.B. co-wrote the manuscript; all authors edited and agreed on the manuscript.

## Acknowledgement

J.M.G. was supported DZD funding and a Marie Curie International Re-integration grant.

## Materials and Methods

### Animal models

Experimental procedures involving live animals were carried out in accordance with animal welfare regulations and with approval of the Regierung Oberbayern (Az 55.2-1-54-2532-187-15 and ROB-55.2-2532.Vet_02-14-157) or ut in concordance with the Karolinska Institutet’s guidelines for the care and use of animals in research, and were approved by the institute’s Animal Ethics Committee respectively. B6 albino and C57BL/6J mice were obtained from Charles River (Germany). *Bbs4^−/−^* mice were generated in the Lupski lab and backcrossed to C57BL/6J^37^. Heterozygous offspring were crossed to produce homozygous *Bbs4^−/−^* and littermate control. *Pifo^wt/-^* mice were crossed to produced homozygous *Pifo^−/−^* mice and littermate controls. βICKO animals were generated by crossing *Pdx1-CreER* mice^38^ to *Ift88^loxP/loxP^* mice^39^ (Volta et al.).

### Islet isolation and transplantation

Islets were isolated by cannulation of the common bile duct of donor animals and infusion of 2.5ml collagenase P, 1.0 mg/ml in HBSS containing 25mM HEPES and 0.2% BSA (Sigma-Aldrich, USA). Inflated pancreata were dissected out and digested at 37°C for 10 min, and cultured in RPMI Medium 1640 (11mM D-glucose) supplemented with 10% fetal bovine serum, 100 IU/ml penicillin, 100 μg/ml streptomycin and 2 mM L-Glutamine (all from Thermo Fisher Scientific, USA). Recipient mice were anesthetized with isoflurane (Baxter, USA) and fixed with a custom-made head holder (Narishige, Japan) as described previously^19^. A small incision was made in the cornea with a 25 G needle and a glass cannula containing islets was inserted through the opening into the anterior chamber of the eye. Around 5-8 islets were carefully positioned on the iris around the pupil. Post-operative analgesia was provided by subcutaneous injection of 2 μg Temgesic (RB Pharmaceuticals Limited, UK).

### Streptozotocin treatment and glucose tolerance test

For Streptozotocin (STZ, Sigma-Aldrich, USA) treatment, animals were fasted for 6 hours and a single dose of 200□ mg STZ per kg body weight in phosphate buffered saline (PBS, Thermo Fisher Scientific, USA) was injected intraperitoneally. For glucose tolerance tests, animals were fasted for 6 hours and 2 g D-glucose (20% glucose solution) per kg body weight was injected intraperitoneally. Blood glucose values were measured afterwards at specified time points using an Accu-Chek Aviva system (Roche, Switzerland).

### In vivo imaging of islet grafts

Imaging was performed between one to twelve weeks post transplantation by confocal microscopy. Images of islet grafts were obtained by a Leica SP5 system with 25× objective (N.A.0.95, Leica Microsystems, Germany). Viscotears (Laboratoires Théa, France) was used as an immersion medium between the lens and the mouse eyes. For visualization of islet vascularization, animals were anesthetized with isoflurane (Baxter, USA). 100□μl of PBS solution containing 5□mg/ml of 70-kDa Texas Red-conjugated dextran (Thermo Fisher Scientific, USA) was injected intravenously prior to imaging. Z-stacks of 2□μm thickness were acquired of every islet graft (Ex.: 561 & 594nm, Em.: 558-564 nm & 600-700 nm). For quantification of vascular leakage, animals were anesthetized with Hypnorm and midazolam (Roche, Switzerland). 100□μl of PBS solution containing 2.5□mg/ml of 40-kDa FITC-conjugated dextran (Thermo Fisher Scientific, USA) was injected intravenously prior to imaging. Time series of z-stacks (2□μm) were acquired every second (Ex.: 488 & 633nm, Em.: 500-550 nm & 630-636 nm). For monitoring the uptake of glucose analogue, animals were anesthetized with Hypnorm and midazolam (Roche, Switzerland). 2-(N-(7-Nitrobenz-2-oxa-1,3-diazol-4-yl)Amino)-2-Deoxyglucose (2-NBDG, Thermo Fisher Scientific, USA) was dissolved in PBS at 5 mg/ml. Recipient mice were anesthetized and a tail vein catheter containing 50 μl 2-NBDG solution was installed. Time series of z-stacks (2□μm) were acquired every second (Ex.: 488 & 633nm, Em.: 500-600nm & 630-636nm) and 2-NBDG solution was injected during imaging.

### Image analysis

Graft volume was estimated by backscatter signal using Volocity image analysis software (Perkin Elmer, USA). Dextran-labeled islet capillaries were identified at different time points, and their diameters and volumes were calculated using Volocity. The same thresholds were used for all groups at all times, and structures smaller than 10μm^3^ were automatically excluded. Re-vascularization rate at one to eight weeks post transplantation was calculated as the ratio of vascular volume at each individual time point and the vascular volume of the same islet at twelve weeks. Vascular density was calculated by counting the volume of islet grafts at twelve weeks and dividing by graft volume. Time lapse confocal image stacks were analyzed and fluorescence intensity was quantified using Fiji^40^ with the plugin Time Series Analyzer V3. Image analysis for histological sections were also performed with Volocity. The region of interest per stack (ROI) was selected based on insulin staining. The surface area and volume of PECAM-1, NG2, and insulin-positive structures were calculated. Islet capillary density was obtained by counting the PECAM-1-positive volume and dividing by the insulin-positive volume enclosed in this region. Pericyte coverage was calculated by counting NG2-positive surface area and dividing by PECAM-1-positive surface area in the same insulin-positive region.

### Cell culture

HEK293FT cells were cultured in Dulbecco’s Modified Eagle’s Medium (DMEM) supplemented with 2 mM L-Glutamine and 10% fetal bovine serum (all from Thermo fisher Scientific, USA). MS1 cells were cultured in Dulbecco’s Modified Eagle’s Medium (DMEM) supplemented with 2 mM L-Glutamine and 5% fetal bovine serum. Human Dermal Microvascular Endothelial Cells (HDMECs) were cultured in Endothelial Cell Growth Medium MV2 with Endothelial Cell Growth Supplement (Promocell, Germany).

### Construction of lentiviral vectors

We used a commercial lentiviral system from Sigma-Aldrich for designing and construct shRNA containing lentiviral vectors. Two target sequences per gene of interest *(Mus musculus)* were selected from MISSIONshRNA database (https://www.sigmaaldrich.com/life-science/functional-genomics-and-rnai/shrna/individual-genes.html). For *Bbs4:* 5’-CCTTGTATTAAGAACCTAG-3’ (shBbs4-1) and 5’-GCATGACCTGACTTACATAAT-3’ (shBbs4-2). For *Ift88*: 5’-TC AGATGCC ATCAACTCATTT-3’ (shIft88-1) and 5’-CTATGAGTCATACAGGTATTT-3’ (shIft88-2). For human *Bbs4:* 5’-TAGTCCTCAGAGTGCTGATAA-3’. Control scrambled sequence: 5’-CCTAAGGTTAAGTCGCCCTCG-3’. The corresponding oligonucleotides were cloned into pLKO.1-puro vector (Sigma-Aldrich, USA) according to manufacturer’s instructions. The efficacy was assessed by relative level of message RNA by qPCR. For reasons of clarity, we refer to more effective shRNA construct shBbs-2 as shBbs4 and shIft88-1 as shIft88 in the text.

### Lentiviral production and endothelial cell transduction

Lentivirus was produced in 293FT cells and harvested following the manufacturer’s guidelines (https://www.sigmaaldrich.com/life-science/functional-genomics-and-rnai/shrna/trc-shrna-products/lentiviral-packaging-mix.html). 1×10^6^ MS1 or 2×10^5^ HDMEC cells were plated in 25 cm^2^ flasks and 1 ml of crude virus containing medium was used for transduction. Transduced cells were further screened in 1.0 μg/ml puromycin (Sigma-Aldrich, USA) for a week before use.

### VEGF-A uptake

5×10^5^ transduced MS1 cells were plated on coverslips in 6 well plates. Confluent cells were starved in DMEM supplemented with 0.5% BSA for 6 hours, and incubated with 50 ng/ml biotinylated mouse VEGF-A (R&D Systems, USA) for 30 min at 37°C. Treated cells were subsequently fixed in 4% paraformaldehyde solution at room temperature for 15 min and permeabilized in 0.1% Triton-X (Sigma-Aldrich, USA) in PBS for 10 min. Fixed cells were then washed 3 times in PBS and blocked with 2% BSA in PBS for 1 hour. To detect the internalized biotinylated VEGF-A, cells were further incubated with Streptavidin-FITC conjugate (1:1000, Thermo Fisher Scientific, USA) at room temperature for 1 hour.

### VEGFR-2 internalization

5×10^5^ transduced HDMEC cells were plated on coverslips in 6 well plates. Confluent cells were starved overnight in DMEM supplemented with 1% BSA. Starved cells were gently washed twice with PBS, and stimulated with 50 ng/ml human VEGF-A (Peprotech, Sweden) for 10 min at 37°C. VEGF-A containing medium was quickly removed and cells were washed twice in ice-cold PBS. Subsequently, cells were fixed in 4% paraformaldehyde solution at room temperature for 15 min, and blocked with 2% BSA in PBS for 1 hour without permeabilization. Further incubation steps with primary and secondary antibodies were described as below.

### VEGF-A ELISA

Isolated islets from control and *Bbs4^−/−^* animals were cultured overnight. 30 islets from each group were picked into 4 well plates containing 450 μl culture medium per well. 72 hours later, supernatants were collected and stored at −80°C. Islets were collected and lysed in 50 μl M-PER buffer (Thermo Fisher Scientific, USA), for quantification of DNA contents by Quant-iT PicoGreen dsDNA Assay Kit (Thermo Fisher Scientific, USA) later according to manufacturer’s instructions. Secreted VEGF-A in the supernatants was measured with mouse VEGF Quantikine ELISA Kit (R&D Systems, USA) according to manufacturer’s instructions.

### EdU labeling

Cell proliferation was detected with Click-iT™ EdU Alexa Fluor™ 488 Imaging Kit (Thermo Fisher Scientific, USA). Briefly, 2×10^5^ transduced MS1 cells were plated on coverslips in 6 well plates. After attachment, cells were starved overnight in DMEM supplemented with 1% BSA, gently washed twice with PBS, and further incubated with 50 ng/ml mouse VEGF-A (Peprotech, Sweden) and 10 μM EdU for 4 hours. Treated cells were fixed in 4% paraformaldehyde solution at room temperature for 15 min and EdU detection was carried out according to manufacturer’s instructions.

### Endothelial cell migration assay

2.5×10^5^ transduced MS1 cells were plated in 12 well plates. Confluent cells were washed gently with PBS, and incubated with 50 ng/ml VEGF-A in DMEM supplemented with 1% BSA. A scratch was made with 1 ml tip in the middle of each well, and images were acquired for documentation of the initial distances between the separated cells. 18 hours later, cells were fixed in 4% paraformaldehyde solution at room temperature for 15 min, and were photographed again for calculating the distances of migration. Images were acquired with BD Pathway 855 Bioimaging Systems using a Olympus 10x objective (BD, USA).

### Endothelial cell matrigel tube formation assay

6×10^4^ transduced MS1 cells were plated in 12 well plates pre-coated with 200 μl Matrigel Matrix (Corning, USA). 2 hours later, culture medium was replaced by DMEM containing 1% BSA and 50 ng/ml VEGF-A. 6 hours later, cells were fixed in 4% paraformaldehyde solution at room temperature for 30 min, and the extent of network formation was observed and photographed. Images were acquired with BD Pathway 855 Bioimaging Systems using a Olympus 10x objective (BD, USA).

### Transmission electron microscopy (TEM)

Islet grafts were dissected out from euthanized animals which have been transplanted previously, and peri-islet iris tissues were removed carefully. Grafts were fixed in 2.5% glutaraldehyde+1% paraformaldehyde in 0.1□M phosphate buffer, pH 7.4 at 4□°C overnight. Samples were further washed in 0.1□M phosphate buffer, pH 7.4 and post-fixed in 2% osmium tetroxide 0.1□M phosphate buffer, pH 7.4 at 4□°C for two hours, dehydrated in ethanol and acetone, and embedded in LX-112 (Ladd Research Industries, Burlington, USA). Ultra-thin sections of 50-60□nm thick were cut by Leica EM UC 6 (Leica Microsystems, Germany) and contrasted with uranyl acetate, followed by lead citrate treatment and checked in Tecnai 12 Spirit Bio TWIN transmission electron microscope (FEI company, The Netherlands) at 100□kV. Digital images were acquired by Veleta camera (Olympus Soft Imaging Solutions, Germany).

### Pancreatic sections and immunofluorescent staining

Pancreata were swiftly removed from euthanized animals, rinsed in PBS and fixed in 4% paraformaldehyde in PBS at room temperature for 2 hours. Tissues were then washed with PBS twice and transferred stepwise to 10%, 20% and eventually 30% sucrose in PBS. Tissues were then embedded in O.C.T. freezing medium (Thermo Fisher Scientific, USA) at −80 °C and subsequently sectioned into 20 μm thick sections.

Cells and islets were fixed in 4% paraformaldehyde solution at room temperature for 15 min or 1 hour, followed by permeabilization in 0.1% Triton-X in PBS, and blocking with 2% BSA or 10% goat serum in PBS respectively. Samples were incubated with primary antibodies, including anti-PEACAM-1 antibody (goat, 1:400, R&D Systems, USA), anti-NG2 antibody (rabbit, 1:100, Millipore, USA), anti-insulin antibody (guinea pig, 1:1000, Dako, Sweden) and anti-VEGFR2 N-terminal antibody (rabbit, 1:100, R&D Systems, USA), at room temperature for 1 hour or at 4 °C overnight. After washing with PBS, cells and tissues were further incubated with secondary antibodies (goat anti-rabbit IgG H+L Alexa Fluor 633, goat anti mouse IgG H+L Alexa Fluor 488 or 633, goat anti guinea pig IgG H+L Alexa Fluor 561 and donkey-anti goat IgG H+L 488; all at 1:1000, Thermo Fisher Scientific, USA) at room temperature for 1 hour, and mounted with ProLong Antifade mountant with DAPI (Thermo Fisher Scientific, USA).

### Western Blots

HDMEC cells were stimulated by 50 ng/ml VEGF-A for indicated lengths of time and lysed in modified RIPA buffer (150 mM NaCl, 50 mM Tris-HCl pH 7.4, 1% NP-40, 0.1% sodium deoxycholate, 1 mM EDTA, all from Sigma-Aldrich, USA) supplemented with protease and phosphatase inhibitors (Mini Complete/PhosphoStop, Roche Diagnostics, Germany). Protein contents were determined using the BCA Protein Assay Kit (Pierce, USA). Protein extracts were separated on 7.5% stain-free precast gels (Bio-Rad, USA) and transferred to 0.22 μm PVDF membranes (GE Healthcare, USA). Blots were probed with primary antibodies, including phospho-VEGFR2 Y1175 antibody (rabbit, 1:1000), VEGFR2 antibody (rabbit, 1:1000), EphrinB2 antibody (rabbit, 1:500), phospho-PLC*γ*1 Y783 antibody (rabbit, 1:1000), PLCγ1 antibody (rabbit, 1:1000), phospho-Akt Thr308/473 antibody (rabbit, 1:1000), Akt1/2 antibody (rabbit, 1:1000), phospho-p44/42 MAPK Thr202/Tyr204 antibody (rabbit, 1:2000) and Erk1/2 antibody (rabbit, 1:2000; all from Cell Signaling Technology, USA), and a-tubulin antibody (mouse,1:1000, Sigma-Aldrich, USA), and subsequently with secondary anti-mouse/ -rabbit IgG (H+L)-HRP conjugates (1:6000, GE Healthcare, USA). Quantification was carried out on original scanned images using Image Lab software (Bio-Rad, USA) and normalization to total protein.

### Quantitative RT-PCR

Total RNA samples were collected from isolated islets of control and *Bbs4^−/−^* animals, or transduced MS1 cells. Tissue and cell samples were lysed and total RNA was isolated using the GeneJET RNA purification kit (Thermo Fisher Scientific, USA) according to manufacturer’s instructions. Extracted mRNA was transcribed into cDNA using the Maxima First Strand cDNA Synthesis Kit (Thermo Fisher Scientific, USA) following manufacturer’s instructions. *Bbs4, Ift88, VEGF-A, VEGFR2, Ang-1, Ang-2, Tie-1, Tie-2, VHL, EphrinA1, EphrinB2, eNOS* and *Dll4* messages were measured using the SYBR Green Master Mix (Thermo Fisher Scientific, USA) with an QuantStudio™ 5 Real-Time PCR System (Applied Biosystems, USA). Relative gene expression level was quantified against the average of *18S, HMBS* and *TBP* messages, and normalized to those of control cells. All primer sequences are listed in Table 1.

**Table 1.**
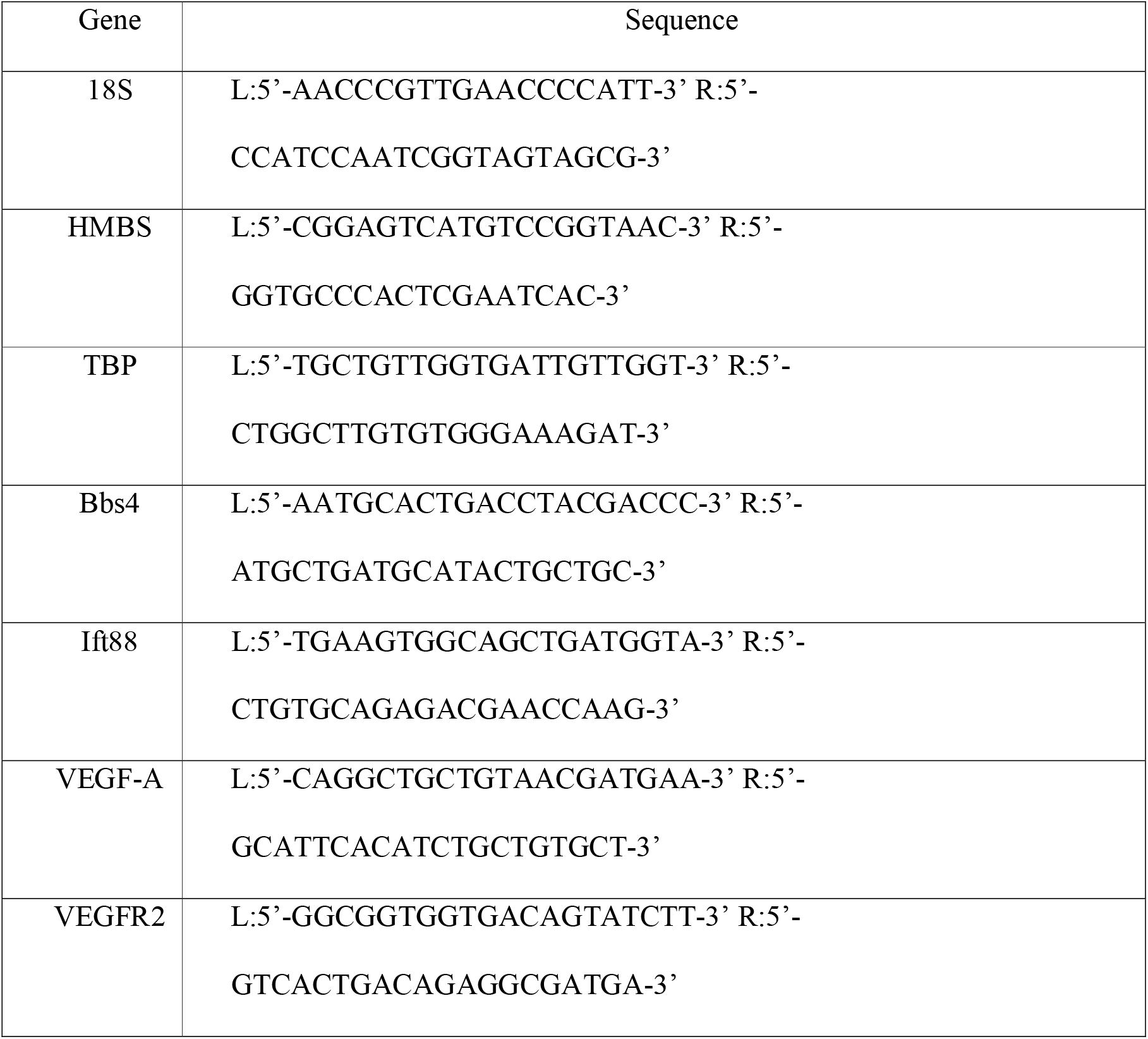
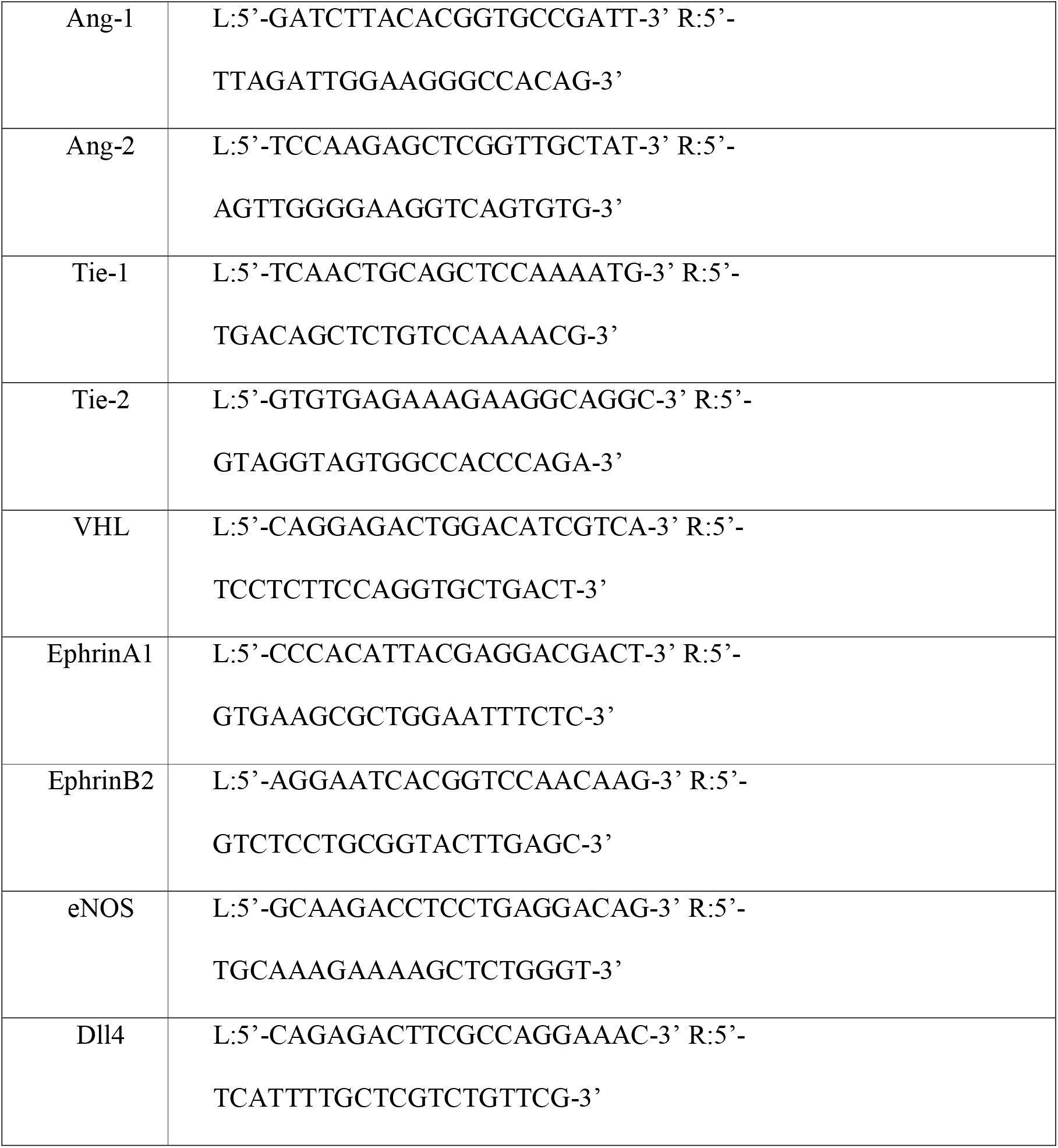
Sequences of qPCR primers

### Statistical tests

All results are presented as either mean ± SEM or shown as box-and-whisker plots (running from minimal to maximal values). Two-way ANOVA test was used for time course analysis, and two-tailed student t-test was used to assess statistical significance for other experiments, with a *p* value < 0.05 considered to indicate significance.

## References

1 Brissova, M. et al. Islet Microenvironment, Modulated by Vascular Endothelial Growth Factor-A Signaling, Promotes ß Cell Regeneration. Cell metabolism 19, 498–511, doi:https://doi.org/10.1016/j.cmet.2014.02.001 (2014).

2 Carmeliet, P. Angiogenesis in health and disease. Nat Med 9, 653–660, doi:10.1038/nm0603-653 (2003).

3 Cleaver, O. & Dor, Y. Vascular instruction of pancreas development. Development 139, 2833–2843, doi:10.1242/dev.065953 (2012).

4 Brissova, M. et al. Intraislet Endothelial Cells Contribute to Revascularization of Transplanted Pancreatic Islets. Diabetes 53, 1318–1325, doi:10.2337/diabetes.53.5.1318 (2004).

5 Brissova, M. et al. Pancreatic Islet Production of Vascular Endothelial Growth Factor-A Is Essential for Islet Vascularization, Revascularization, and Function. Diabetes 55, 2974–2985, doi:10.2337/db06-0690 (2006).

6 Vaithilingam, V., Sundaram, G. & Tuch, B. Islet cell transplantation. Curr Opin Organ Transpl 13, 633–638, doi:10.1097/MOT.0b013e328317a48b (2008).

7 Christofori, G., Naik, P. & Hanahan, D. Vascular endothelial growth factor and its receptors, flt-1 and flk-1, are expressed in normal pancreatic islets and throughout islet cell tumorigenesis. Mol Endocrinol 9, 1760–1770, doi:10.1210/mend.9.12.8614412 (1995).

8 Reinert, R. B. et al. Vascular endothelial growth factor-a and islet vascularization are necessary in developing, but not adult, pancreatic islets. Diabetes 62, 4154–4164, doi:10.2337/db13-0071 (2013).

9 Wheatley, D. N., Wang, A. M. & Strugnell, G. E. Expression of primary cilia in mammalian cells. Cell biology international 20, 73–81, doi:10.1006/cbir.1996.0011 (1996).

10 Nauli, S. M., Jin, X. & Hierck, B. P. The mechanosensory role of primary cilia in vascular hypertension. Int J Vasc Med 2011, 376281, doi:10.1155/2011/376281 (2011).

11 Kallakuri, S. et al. Endothelial cilia are essential for developmental vascular integrity in zebrafish. J Am Soc Nephrol 26, 864–875, doi:10.1681/ASN.2013121314 (2015).

12 Dinsmore, C. & Reiter, J. F. Endothelial primary cilia inhibit atherosclerosis. EMBO reports, doi:10.15252/embr.201541019 (2016).

13 Vion, A. C. et al. Primary cilia sensitize endothelial cells to BMP and prevent excessive vascular regression. J Cell Biol 217, 1651–1665, doi:10.1083/jcb.201706151 (2018).

14 Oh, E. C., Vasanth, S. & Katsanis, N. Metabolic regulation and energy homeostasis through the primary Cilium. Cell Metab 21, 21–31, doi:10.1016/j.cmet.2014.11.019 (2015).

15 Volta, F. & Gerdes, J. M. The role of primary cilia in obesity and diabetes. Ann N Y Acad Sci 1391, 71–84, doi:10.1111/nyas.13216 (2017).

16 Gerdes, J. et al. Ciliary dysfunction impairs beta-cell insulin secretion and promotes development of Type 2 Diabetes in rodents. Nature communications 5, 5308, doi:10.1038/ncomms6308 (2014)(2014).

17 Konishi, M. et al. Endothelial insulin receptors differentially control insulin signaling kinetics in peripheral tissues and brain of mice. Proc Natl Acad Sci U S A 114, E8478–E8487, doi:10.1073/pnas.1710625114 (2017).

18 Goligorsky, M. S. Vascular endothelium in diabetes. Am J Physiol Renal Physiol 312, F266–F275, doi:10.1152/ajprenal.00473.2016 (2017).

19 Speier, S. et al. Noninvasive high-resolution in vivo imaging of cell biology in the anterior chamber of the mouse eye. Nat Protoc 3, 1278–1286, doi:10.1038/nprot.2008.118 (2008).

20 Linn, T. et al. Angiogenic capacity of endothelial cells in islets of Langerhans. FASEBjournal: official publication of the Federation of American Societies for Experimental Biology 17, 881–883, doi:10.1096/fj.02-0615fje (2003).

21 Nyqvist, D., Köhler, M., Wahlstedt, H. & Berggren, P.-O. Donor Islet Endothelial Cells Participate in Formation of Functional Vessels Within Pancreatic Islet Grafts. Diabetes 54, 2287–2293, doi:10.2337/diabetes.54.8.2287 (2005).

22 Ilegems, E. et al. Light scattering as an intrinsic indicator for pancreatic islet cell mass and secretion. Sci Rep 5, 10740, doi:10.1038/srep10740 (2015).

23 Henderson, J. R. & Moss, M. C. A morphometric study of the endocrine and exocrine capillaries of the pancreas. Q J Exp Physiol 70, 347–356 (1985).

24 Rodriguez-Diaz, R. et al. Paracrine Interactions within the Pancreatic Islet Determine the Glycemic Set Point. Cell Metab 27, 549–558 e544, doi:10.1016/j.cmet.2018.01.015 (2018).

25 Esser, S. et al. Vascular endothelial growth factor induces endothelial fenestrations in vitro. J Cell Biol 140, 947–959 (1998).

26 Roberts, W. G. & Palade, G. E. Increased microvascular permeability and endothelial fenestration induced by vascular endothelial growth factor. J Cell Sci 108 (Pt 6), 2369–2379 (1995).

27 Lammert, E. et al. Role of VEGF-A in vascularization of pancreatic islets. Curr Biol 13, 1070–1074 (2003).

28 Sawamiphak, S. et al. Ephrin-B2 regulates VEGFR2 function in developmental and tumour angiogenesis. Nature 465, 487–491, doi:10.1038/nature08995 (2010).

29 Wang, Y. et al. Ephrin-B2 controls VEGF-induced angiogenesis and lymphangiogenesis. Nature 465, 483–486, doi:10.1038/nature09002 (2010).

30 Nakayama, M. et al. Spatial regulation of VEGF receptor endocytosis in angiogenesis. Nat Cell Biol 15, 249–260, doi:10.1038/ncb2679 (2013).

31 Bruns, A. F. et al. Ligand-stimulated VEGFR2 signaling is regulated by co-ordinated trafficking and proteolysis. Traffic 11, 161–174, doi:10.1111/j.1600-0854.2009.01001.x (2010).

32 Pi, X., Xie, L. & Patterson, C. Emerging Roles of Vascular Endothelium in Metabolic Homeostasis. Circ Res 123, 477–494, doi:10.1161/CIRCRESAHA.118.313237 (2018).

33 Iwashita, N. et al. Impaired insulin secretion in vivo but enhanced insulin secretion from isolated islets in pancreatic beta cell-specific vascular endothelial growth factor-A knock-out mice. Diabetologia 50, 380–389, doi:10.1007/s00125-006-0512-0 (2007).

34 Honkura, N. et al. Intravital imaging-based analysis tools for vessel identification and assessment of concurrent dynamic vascular events. Nat Commun 9, 2746, doi:10.1038/s41467-018-04929-8 (2018).

35 Wang, Y., Jin, Y., Lavina, B. & Jakobsson, L. Characterization of multi-cellular dynamics of angiogenesis and vascular remodelling by intravital imaging of the wounded mouse cornea. Sci Rep 8, 10672, doi:10.1038/s41598-018-28770-7 (2018).

36 Buchholz, B. M. et al. Revascularization Time in Liver Transplantation: Independent Prediction of Inferior Short-and Long-term Outcomes by Prolonged Graft Implantation. Transplantation 102, 2038–2055, doi:10.1097/TP.0000000000002263 (2018).

37 Eichers, E. et al. Phenotypic characterization of Bbs4 null mice reveals age-dependent penetrance and variable expressivity. Human genetics 120, 211–226 (2006).

38 Zhang, H., Fujitani, Y., Wright, C. & Gannon, M. Efficient recombination in pancreatic islets by a tamoxifen-inducible Cre-recombinase. Genesis 42, 210–217 (2005).

39 Davenport, J. et al. Disruption of Intraflagellar Transport in Adult Mice leads to Obesity and Slow-Onset Cystic Kidney Disease. Current biology: CB 17, 1586–1594 (2007).

40 Schindelin, J. et al. Fiji: an open-source platform for biological-image analysis. Nat Methods 9, 676–682, doi:10.1038/nmeth.2019 (2012).

